# Genome-wide scans of selection highlight the impact sof biotic and abiotic constraints in natural populations of the model grass *Brachypodium distachyon*

**DOI:** 10.1101/246090

**Authors:** Yann Bourgeois, Christoph Stritt, Jean-Claude Walser, Sean P. Gordon, John P. Vogel, Anne C. Roulin

## Abstract

Grasses are essential plants for ecosystem functioning. Quantifying the selective pressures that act on natural variation in grass species is therefore essential regarding biodiversity maintenance. In this study, we investigate the selection pressures that act on two distinct populations of the grass model *Brachypodium distachyon* without prior knowledge about the traits under selection. We took advantage of whole-genome sequencing data produced for 44 natural accessions of *B. distachyon* and used complementary genome-wide scans of selection (GWSS) methods to detect genomic regions under balancing and positive selection. We show that selection is shaping genetic diversity at multiple temporal and spatial scales in this species and affects different genomic regions across the two populations. Gene Ontology annotation of candidate genes reveals that pathogens may constitute important factors of positive and balancing selection in *Brachypodium distachyon*. We eventually cross-validated our results with QTL data available for leaf-rust resistance in this species and demonstrate that, when paired with classical trait mapping, GWSS can help pinpointing candidate genes for further molecular validation. Thanks to a near-base perfect reference genome and the large collection of freely available natural accessions collected across its natural range, *B. distachyon* appears as a prime system for studies in ecology, population genomics and evolutionary biology.

## Introduction

Grasses cover more than 40% of the world land area (Gibson, 2009) and dominate a wide variety of ecosystems, from tropical to temperate regions (Clayton, 1981; Gibson, 2009). Grasses also play a key role in eco-and agrosystem functioning as they provide habitats for many animal species (Groves, 2000) and represent the main source of grain and forage (Stromberg 2011). Increasing crop production to meet the food and energy requirements of the world’s growing population is, however, putting great pressure on natural grasslands (Wallace, 1997; Helm *et al.*, 2009; Ceballos *et al.*, 2010). Faced with constant deterioration and fragmentation due to human activities (Kiviniemi, 2002), these ecosystems are highly endangered (Ceballos *et al.*, 2010), but little is known about their evolutionary resilience. Assessing the genetic basis of adaptation and quantifying the selection pressures that act on natural variation in grass species is therefore crucial with respect to biodiversity maintenance and food security.

To date, reciprocal transplant experiments have been extensively used to test for adaptive differentiation across populations (for review see Savolainen *et al.*, 2013). Based on a “home vs. foreign” effect on fitness, reciprocal transplants are indeed powerful to unravel overall genotype by environment (GxE) interactions and demonstrated the prevalence of local adaptation in grasses and plants in general (for review see Bischoff *et al.*, 2006; Wadgymar *et al.*, 2017). However, reciprocal transplant experiments use information such as survival, vegetative growth or seed production to measure the effect of the habitat on fitness (Bischoff et al. 2006). Hence they provide little insight into the functional and genetic bases of adaptation, unless combined with trait mapping such as quantitative trait locus (QTL) analyses and genome-wide association studies (GWAS) (Latta 2009). QTL analyses and GWAS, on the other hand, are largely constrained by the effort and time required for high-resolution mapping. In grasses, while these trait-by-trait approaches have been valuable to decipher the genetic architecture of important characters with regard to crop genetic improvement (Huang et al. 2002; Barbieri et al. 2012; Morris et al. 2013; Slavov et al. 2014), they remain of limited value to grasp the overall selective forces that act on natural populations.

An efficient alternative to provide insights about evolutionary forces in natural populations consists in identifying genes under selection at a whole genome scale, then describing their function and the type of selection acting on them (Mitchell-Olds et al. 2007). For instance, new mutations that are beneficial in some populations can be positively selected and are more likely to quickly increase in frequency. Such so-called selective sweeps tend to reduce genetic diversity, increase differentiation among populations, and lead to extended haplotypes in the vicinity of the locus under selection due to genetic hitchhiking (Nielsen 2005; Hermisson 2009). Various genome-wide selection scans (GWSS) methods have been developed to detect such footprints of positive selection (Tang et al. 2007; Gautier et al. 2012; Stamatakis et al. 2013; Messer 2015) and GWSS are now emerging as complementary approaches to classical trait mapping.

A first class of GWSS is based on cross-population comparison. These methods scan the genome by comparing the lengths of haplotypes across populations and are powerful to identify hard selective sweeps (Tang *et al.*, 2007), i.e. regions in the genome where one haplotype has undergone recent and near-complete fixation in one of the studied populations in response to selection (Figure 1A). Yet, if the advantageous mutation has been present in the population for long enough and recombined before selection took place, then several different haplotypes may increase in frequency (Figure 1B). Such events (“soft sweeps”), as well as partial hard sweeps, are less efficiently detected by GWSS based on cross-population comparison (Schlamp et al. 2016) since they do not erase polymorphism to the same extent as completed hard sweeps. Soft-sweeps can be yet detected by a second class of GWSS which are also based on extended haplotypes but are applied at the within population level. These scans, by measuring mean pairwise differences in haplotype length along the genome in a given population, can indeed identify loci under selection independently of the number of haplotypes (Figure 1B) or of the completeness of the sweep (Schlamp et al. 2016). Therefore, within-population GWSS can detect not only soft-sweeps, but also mutations that are undergoing selection and did not reach complete fixation within a population.

**Figure 1:**
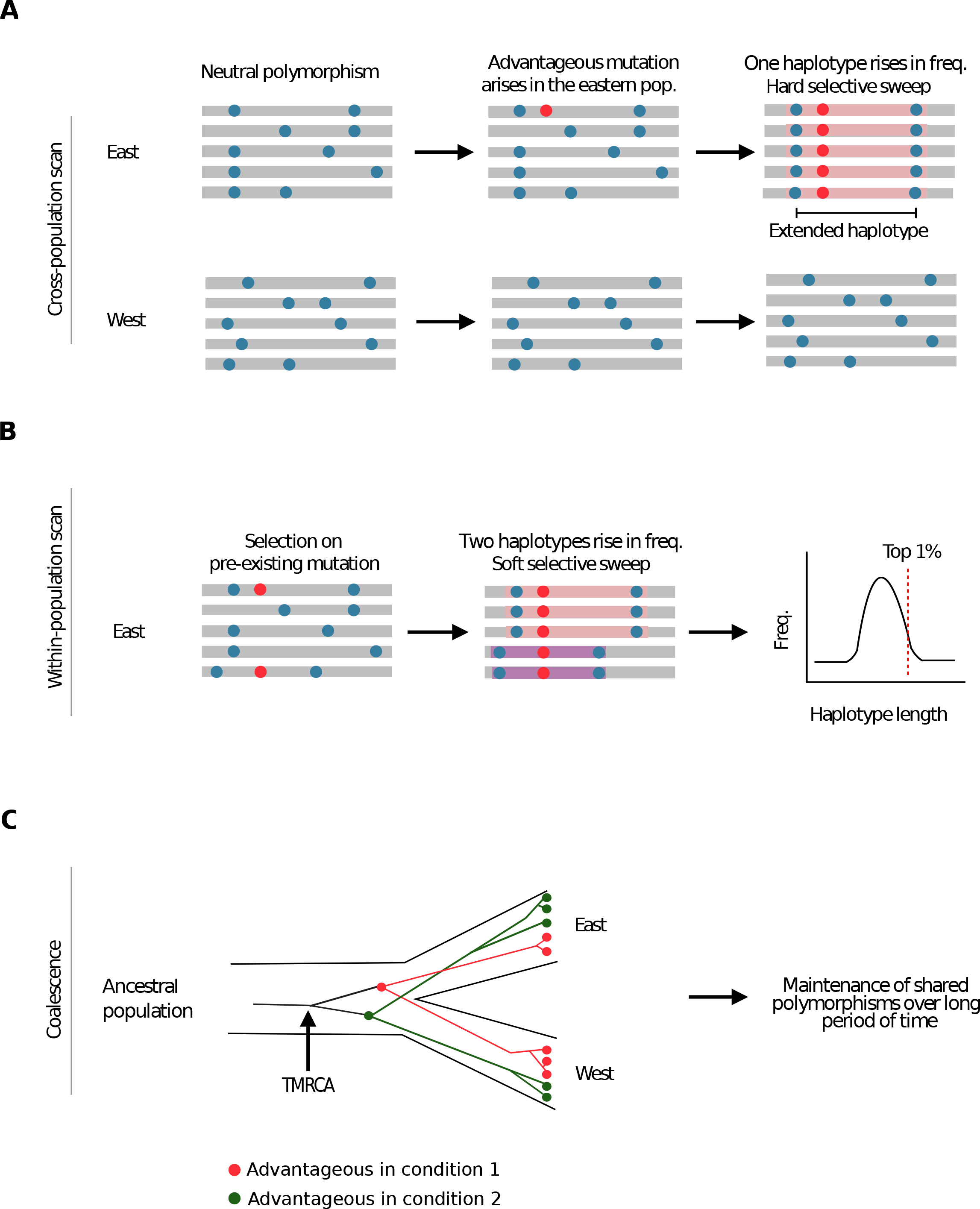
Schematic representation of the regimes of selection analyzed in this study. A) Cross-population tests: in case of positive selection on a new mutation, SNPs linked to the advantageous mutation will also rise in frequency, leading to extended haplotypes in the vicinity of the selected mutation. Cross-population tests detect haplotypes that are positively selected in one population by estimating the length of haplotypes around each allele at a core SNP, then comparing these lengths between populations. Such tests are designed to identify hard-sweep, i.e. region where one haplotype reached near fixation B) Within-population tests: these scans measure haplotype length at a focal locus and compare it to the average length of haplotypes along the genome in a given population. They allow recovering regions with haplotypes longer than average (top 1% outliers) and can identify loci under selection independently of the number of haplotypes or of the completeness of the sweep C) Coalescence approach: balancing selection can maintain shared polymorphisms across populations due the habitat heterogeneity or negative frequency-dependent selection. Associated to old alleles, long-term balancing selection can be detected by calculating time to most recent common ancestor (TMRCA), which should increase at loci maintained by selection over long period of time.

Scans for positive selection are nonetheless not exempt of shortcomings and, as any computational approach, can lead to the detection of false positives. First, these tests often rely on the assumption that regions above a given threshold for the test statistics should be enriched in selected genes. However, defining which threshold should limit the amount of false positives is not straightforward (Pavlidis et al. 2012). In addition, demographic events such as bottleneck may also lead to the fixation of long haplotypes and mimics signal of selection (Schrider & Kern, 2016). This is especially true when the bottleneck is recent and that recombination did not have enough time to break haplotype blocks. While some GWSS are now able to incorporate bottlenecks or expansions in their models (Schrider & Kern, 2016), disentangling the effects of selection from demography remains notoriously challenging as it implies to precisely model population size evolution over time. Comparing and validating cross-and within-population GWSS with approaches that take into account the demographic history of each population may therefore constitute the best way to identify genes under different regimes of positive selection with high confidence.

Furthermore, while local adaptation is commonly associated to positive selection on new advantageous polymorphisms, recent studies have demonstrated that balancing selection is also playing an important role in this evolutionary process (Mitchell-Olds et al. 2007; Rasmussen et al. 2014; Wu et al. 2017). The term balancing selection is an “umbrella” concept (Fijarczyk & Babik, 2015) which describes the maintenance of genetic diversity over longer periods of time through adaptation to spatial heterogeneity, heterozygote advantage and negative frequency-dependent selection (Mitchell-Olds et al. 2007; Rasmussen et al. 2014). This process is more difficult to detect than positive selection (Fijarczyk & Babik, 2015) since older alleles had more time to recombine and may lead to narrow signatures around selected sites (Teixeira et al. 2014). As a consequence, the effect of balancing selection is still largely overlooked in genome scans, which remain strongly biased towards the detection of recent positive selection (Hassl & Payseur, 2016). It is now possible, however, to model the coalescent process along chromosomes and to identify genomic regions with unusually long times to the most recent common ancestor (TMRCA, Figure 1C), a signature which can arise through balancing selection (Rasmussen et al. 2014).

In this study, we capitalize on the near base-perfect quality of the reference genome of the Mediterranean grass *Brachypodium distachyon* (https://phytozome.jgi.doe.gov) to investigate how both positive and balancing selection are shaping diversity in this species. In the last decade, *B. distachyon* has been developed as a powerful model for research on temperate grass species as it is closely related to major crop cereals and to some of the grasses used for biofuel production (The International Brachypodium Consortium 2010). Entirely sequenced, its small diploid genome (272Mb) is fully assembled into five chromosomes and has been exhaustively annotated (The International Brachypodium Consortium 2010). In addition, *B. distachyon* is broadly distributed around the Mediterranean rim (Dell’Acqua *et al.*, 2014; Gordon *et al.*, 2014; Tyler *et al.*, 2016), providing access to natural populations from contrasting habitats for which a large collection has been collected. It constitutes therefore a unique and prime system to investigate the genetic basis of local adaptation in natural grass populations, opening the way to further fundamental and applied research.

We took advantage of whole-genome sequencing data produced for 44 *B. distachyon* natural accessions originating mainly from Spain and Turkey (Gordon *et al.*, 2017). Those accessions are genetically distinct, occur in the eastern and western outer range of the species distribution and were selected to maximize genetic differentiation and identify regional selective pressures that may have been important during recolonization following the last glacial maximum (Stritt et al. 2018). Making use of the 6 million SNPs we identified in these genomes, we combined a cross-population (Rsb; Tang *et al.*, 2007) and a within-population scan (H-scan; Schlamp *et al.*, 2016) to an integrated demography approach (diploS/HIC; Kern & Schrider, 2018) to detect high-confidence genomic regions under different regimes of positive selection. We also tested for regions under balancing selection with the program ARGweaver (Rasmussen et al. 2014) and asked: i) At what time and geographical scale is selection acting in *B. distachyon* populations? ii) What are the selective constrains that shape diversity and adaptation in these populations? iii) Can we recover candidates for local adaptation identified from QTL analyses performed in controlled conditions?

## Results

### Population structure and genetic diversity

In this study, we used whole-genome sequencing data (paired-end; Illumina technology) with a 86-fold median coverage of 44 *B. distachyion* accessions originating from Turkey, Iraq, Spain and France (Figure 2A, Table S1, Gordon *et al.*, 2017). After filtering, we identified 6,204,029 SNPs. An Admixture analysis, where K=2 was identified as the best model (Figure S1), highlighted two distinct genetic clusters, an eastern and a western one, with extremely little admixture between the two (Figure 2B). For the rest of the study, and even though these accessions belong to two genetic groups rather than *stricto sensu* populations, accessions from Turkey and Iraq will be referred to as the eastern population while accessions from Spain and France will be referred to as the western population.

**Figure 2:**
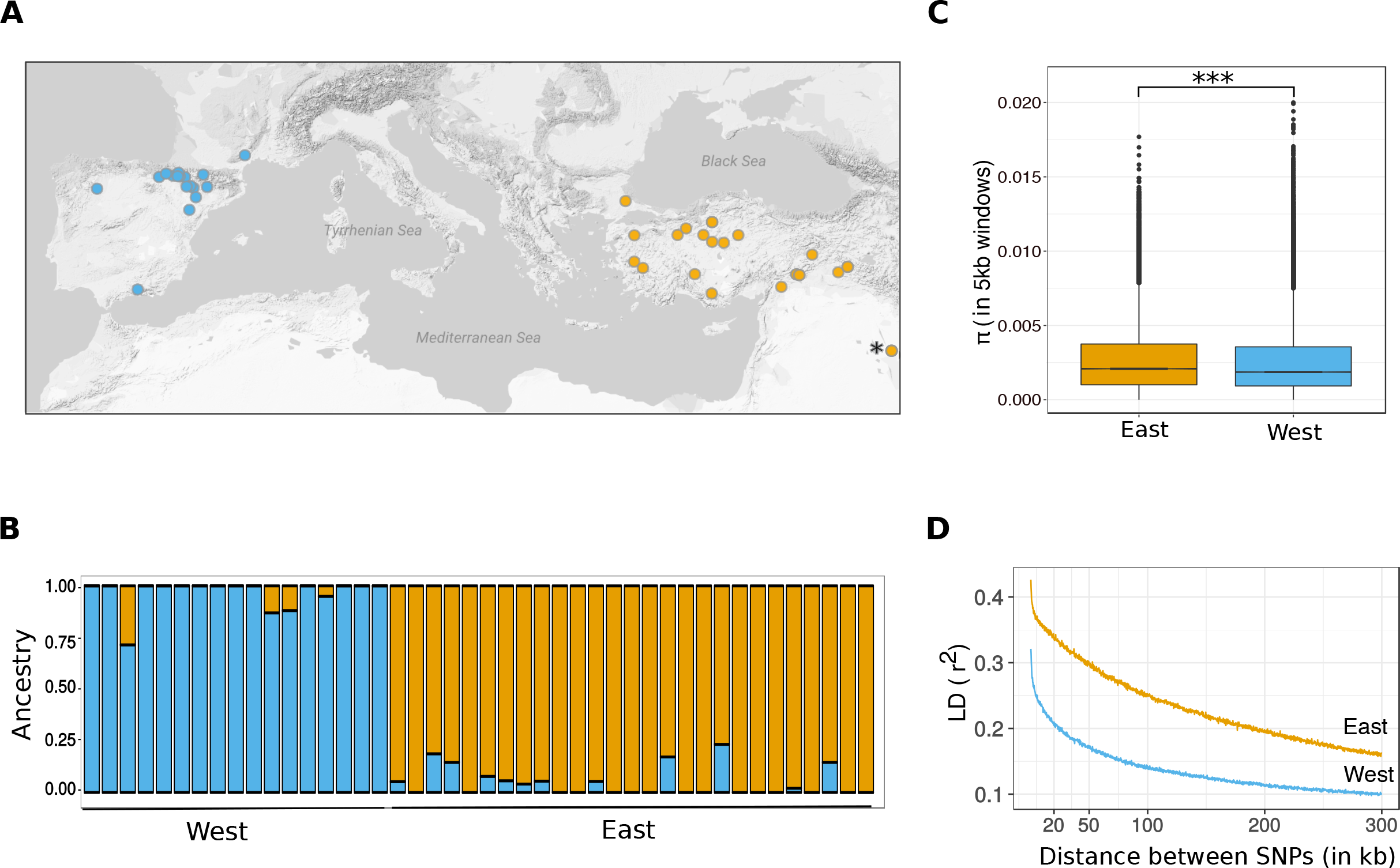
Geographical origin and genetic diversity of the 44 accessions. A) Geographical origin of the 44 accessions used in the study. The asterisk indicates the origin of the reference accession Bd21 B) Structure plot (K=2) displaying the limited level of admixture between the eastern and western population. Each bar represents the genetic data of one accession. An accession presents some admixture when the bar displays different colors. The height of the bar is proportional to the admixture level C) Nucleotide diversity (π in 5 kb windows) within the eastern and western populations D) LD decay in the eastern and western populations.

The western population showed a lower level of nucleotide diversity (median π value in east = 0.0021, median π value in west = 0.00186; Wilcoxon test; P-value < 2.2e-16, Figure 2C) than the eastern one. Excluding the reference accession Bd21, which has been artificially inbred before sequencing, the level of heterozygosity in these accessions ranges from 4 to 17.4% (Table S1) but median values (7.8 and 6.3 in the eastern and western population respectively) did not statistically differ between the two populations (Wilcoxon test; P-value = 0.31). Eventually, linkage disequilibrium (LD) decays faster in the western population than in the eastern one (Figure 2 D).

### Functional clustering of the genome of B. distachyon and subsequent filtering

To assess whether the genome of *B. distachyon* harbors functional clusters of genes, we first performed a GO annotation for the 32,712 genes annotated in the reference genome. We then controlled for potential gene clustering by following the procedure described in (Al-Shahrour *et al.*, 2010). Briefly, we split the genome into overlapping windows of 50 consecutive genes and performed enrichment analyses on each window. We identified 272 windows significantly enriched for at least one biological process (Table S2). Several windows were enriched for processes that may be associated to adaptation to local environmental conditions such response to stress and defense response (Table S2).

A classical approach applied to analyze the results of GWSS consists in selecting genomic regions containing top 1% outliers for signals of selection and then assessing whether some biological functions or processes are significantly over-represented in these regions through a Gene Ontology (GO) annotation. We anticipated that the nonrandom organization of *B. distachyon* genome we observed could lead to an over representation of some biological pathways or functions in small genomic regions and to an artificial enrichment for some GO terms in GWSS 1% outlier regions (Pavlidis et al. 2012). This prompted us to narrow down the top 1% outlier regions by keeping only the genes located at and in the immediate vicinity (−10% of the peak value) of each of the peaks of selection for the cross-(Rsb test) and within-(H-scan) population scans of selection presented in the following section. In contrast, the outputs of the coalescence and demography approaches, which are both window-based (see methods), were not filtered.

### Detection of hard selective sweeps through cross-population contrasts

When new mutations providing a selective advantage rise in frequency through positive selection, neutral mutations that are physically close tend to remain strongly linked to them. Recent positive selection should therefore lead to a signature of long haplotypes near selected mutations. We used the Rsb test (see methods) to contrast the two populations (Tang et al. 2007) and to detect regions where one haplotype (hard sweep, Figure 1A) reached near or complete fixation a recent past in one of the two populations. By doing so, we identified 312 regions harboring 824 genes and 319 regions harboring 1212 genes potentially under selection in the eastern and western populations respectively.

### Detection of soft or ongoing selective sweeps through a within-population scan

The Rsb test is less efficient to detect soft-sweeps, i.e. regions where more than one haplotype increased in frequency (Figure 1B), or to identify ongoing selective sweep where not all individuals within the population display haplotype extension in the region under selection. As an alternative, we used the program H-scan (see methods) to detect loci associated to soft selective sweeps or undergoing positive selection within each population. We identified 142 regions harboring 487 genes and 79 regions harboring 463 genes in the eastern and western population respectively. As expected given the specificity of each test, only 49 and 58 genes were common to the H-scan and Rsb approaches in the eastern and the western populations respectively.

### Integration of demography and detection of high-confidence gene sets under positive selection

As presented in the introduction, bottlenecks can mimic signals of selection as they also lead to the fixation of long haplotypes. To alleviate this issue, we applied diploS/HIC, a recently developed machine-learning approach to detect selective sweeps and classify them as hard or soft (Schrider and Kern 2016; Kern and Schrider 2018) while integrating the demographic history of the eastern and western population previously inferred by (Stritt et al. 2018). One of the major advantages of diploS/HIC is that the algorithm is robust to demographic model misspecification (Schrider and Kern 2018). By providing a confusion matrix, it also estimates false positive and false negative rates and allows a better assessment of the confidence levels expected under different scenarios of selection.

We first trained the program to identify regions under positive selection. To do so, genomic windows were simulated under neutrality, with a hard sweep or with a soft sweep (see methods). The algorithm was able to recover from the simulations 71% and 91.5% of hard sweeps in western and eastern populations respectively (Figure S2 for and example of confusion matrix). For soft sweeps, these proportions were of 71% and 83.7%. The proportion of neutral simulations being classified as hard sweeps was low, accounting for 1.25% and 0.04% of the simulations in western and eastern populations. These proportions were higher for soft sweeps, particularly in the western population (11.28%) but still remained low in the eastern population (2.6%). These results suggest a good sensitivity and specificity of diploS/HIC to detect regions that are under selection, especially in the eastern population.

We then ran diploS/HIC on our actual dataset. Windows classified as selective sweeps by diploS/HIC on this dataset recovered 34% (164 genes) and 38% (312 genes) of the genes detected by the H-scan and Rsb tests in the eastern population, as well as 55% (258 genes) and 53% (644 genes) of the genes detected by the H-scan and Rsb tests in the western population. DiploS/HIC recovered more genes detected by the Rsb and H-scan tests than expected by chance (Fisher test, all P-values < 1.8e-12). These four new genes sets (intersects between diploS/HIC-Rsb East, diploS/HIC-Rsb West, diploS/HIC-Hscan East, diploS/HIC-Hscan West) are considered as high-confidence genes under positive selection (Table S3 for a list). Unless mentioned otherwise, those sets will be referred to as genes under positive selection in the discussion.

The high-confidence gene set diploS/HIC-Rsb of the eastern population displayed a significant enrichment for genes involved in stress response and defense response (Table 1). Among other GO term enrichments, the eastern high-confidence gene set diploS/HIC-Hscan was significantly enriched for gene involved in response to cadmium ion (see Table 1 for more significant GO terms). As we observed that some of the well-known types of plant resistance genes (i.e. R-genes harboring NBS-LRR domains) were not associated with the process of defense response in the GO analysis, we specifically counted how many genes containing NBS-LRR domains were present in the four high-confidence gene sets and compared those to the number genes with NBS-LRR domains annotated in the whole genome. Both high-confidence gene sets (diploS/HIC-Hscan and diploS/HIC-Rsb) of the eastern population displayed a significant enrichment for R-genes (P-value = 1.7e-06 and 1.3e-05 respectively). This was not the case for the high-confidence gene sets of the western population (both P-value > 0.6).

**Table 1:** Significant GO terms associated to each high-confidence gene set.

| **Test** | **Significant GO process** | **Pvalue** |
| --- | --- | --- |
| **TMRCA** | Phosphorylation | 3.13E-08 |
| **DiploS/HIC-Rsb East** | Response to stress including:<br>- defense response | 5.00E-04<br>9.4E-04 |
| **DiploS/HIC-Rsb West** | Amide transport | 5.00E-03 |
| **DiploS/HIC-Hscan East** | Response to cadmium ion<br>Small molecule metabolic process | 6.6E-04<br>3.8E-03 |
| **DiploS/HIC-Hscan West** | Pyruvate metabolic process<br>Carbohydrate derivative metabolic process<br>Phospholipid metabolic process | 3.80E-03<br>3.80E-03<br>3.40E-03 |

### Genomic regions affected by positive selection

The regions common to diploS/HIC-Hscan contained slightly more genes in the western than in the eastern population (Wilcoxon test, P-value=0.04; Figure 3A), while there was no significant difference between the sizes of the diploS/HIC-Rsb regions in the eastern and western populations (Wilcoxon test, P-value=0.07; Figure 3A).

**Figure 3:**
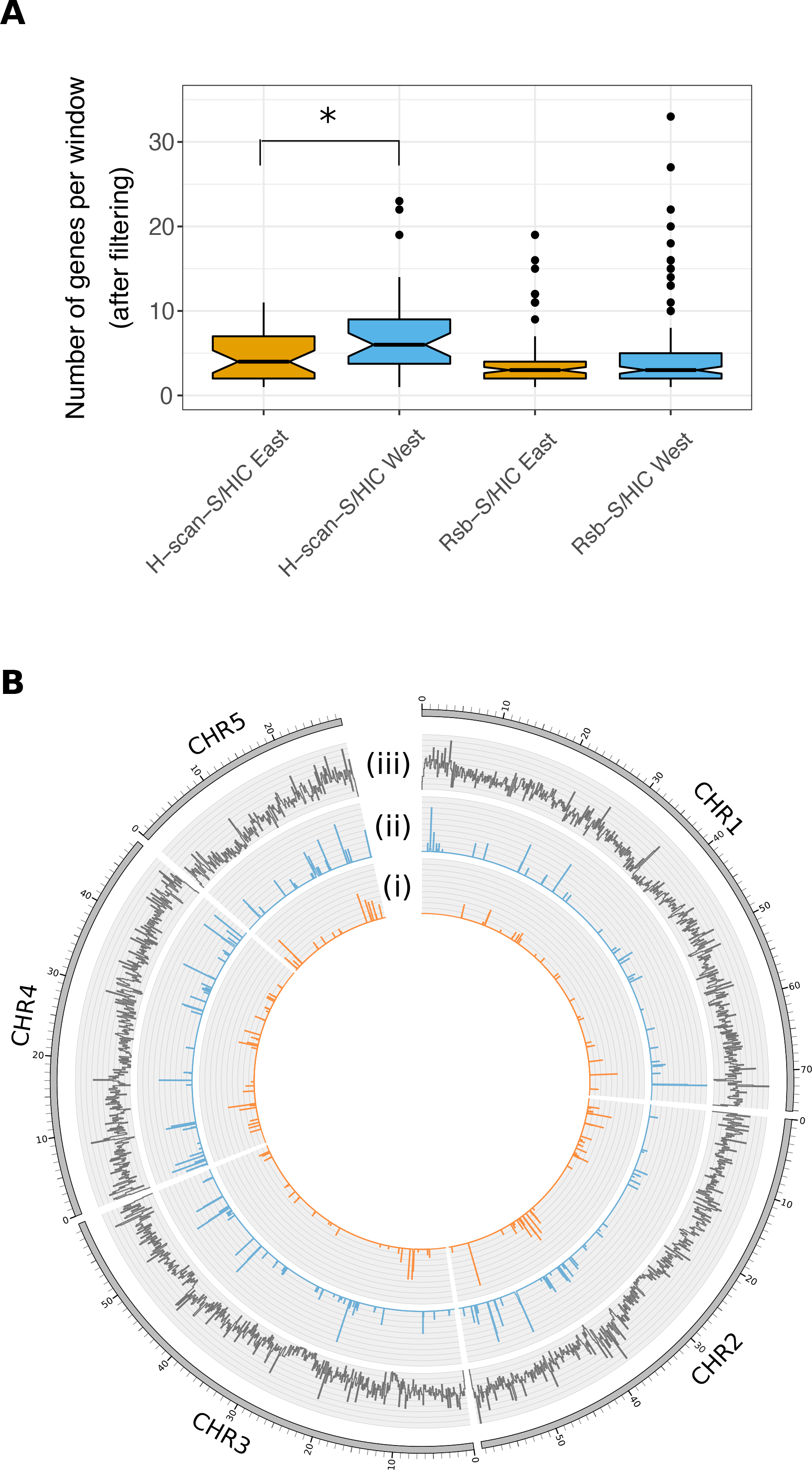
Density of genes under selection. A) Number of genes per high-confidence region for the high-confidence gene sets B) Circular representation of gene density along chromosomes (i) density of genes under positive selection in the eastern population (DiploS/HIC-Rsb and DiploS/HIC-Hscan combined), scaled from 0 to 20 genes per 100 kb (ii) density of genes under positive selection in the western population (DiploS/HIC-Rsb and DiploS/HIC-Hscan combined), scaled from 0 to 20 genes per 100 kb (iii) overall gene density, scaled from 0 to 40 genes per 100 kb.

We also tested whether distinct loci/genomic regions were affected by recent positive selection in the two populations using linear models. While we found a significant association between the density of genes under positive selection (S/HIC-Rsb and S/HIC-Hscan outliers combined) in 100 kb windows in the eastern and western populations on all chromosomes, all R^2^ were inferior to 1.307e-06. This result indicates that positive selection mostly affects distinct loci across the two populations (as depicted on Figure 3B). Interestingly, we observed that some genomic regions harbor higher density of genes under selection than the rest of the genome. Thus, genes under positive selection appear to be non-randomly distributed along chromosomes in the two populations (Figure 3B).

### Genes with extremely long coalescence time

Balancing selection can be detected with coalescence approaches, as ancient alleles are associated with older coalescence times (Charlesworth 2006). We used the software ARGweaver to detect footprints of balancing selection, a method that has robustly identified signatures of balancing selection in recently expanded human populations (Rasmussen et al. 2014). Briefly, ARGweaver models the coalescent process along chromosomes and across non-recombining haplotype blocks to infer their evolutionary history (see methods). It allows recovering several statistics such as times to the most recent common ancestor (TMRCA, Figure 1C), which should be increased near ancient alleles such as those under ancient balancing selection. By doing so, we identified 72 regions harboring 115 genes under what will be referred to as long-term balancing selection in the following (Table S3 for a list). This gene set harbors a significant enrichment for genes involved in phosphorylation (Table 1).

### Identification of candidate genes in known QTL regions

Combining association mapping and analyses of selection constitutes a powerful approach to identify candidate genes and to address their selective regime. We identified many candidate regions harboring resistance genes. As a proof of concept, we therefore aimed at assessing whether regions identified as resistance loci against known pathogens were also highlighted in our selection scans. In *B. distachyon*, the genetic basis of resistance to the rust fungus *Puccinia brachypodii* has been deciphered through a QTL mapping (Barbieri et al. 2012) which showed that leaf-rust resistance is controlled by three main QTL located on chromosome 2 (from nucleotide 37,949,269 to 40,903,216), 3 (from nucleotide 13,943,000 to 14,512,222) and 4 (from nucleotide 9,649,152 to 10,679,750). For the rest of the study, these regions will be referred to as QTL_rust-2_, QTL_rust-3_ and QTL_rust-4_.

We screened these three regions for evidence of selection and found signals of positive selection in QTL_rust-3_ and QTL_rust-4_. In QTL_rust-3_, we found that one gene (Bradi3g16331: 14,502,928-14,503,290) belonged to the eastern high-confidence gene set diploS/HIC-Rsb (Figure 4A, left panel). This gene is located 60 kb upstream the QTL peak identified in this region (Barbieri et al. 2012) and codes for an unknown protein. It is flanking a serine/threonine phosphatase (Bradi3g16320: 14,486,831-14,488,838; Figure 4A, left panel) which was also identified as being under selection in the eastern population with the Rsb test. The latter gene was however not recovered by diploS/HIC but classified as a locus being linked to a selected site (see methods). This 16kb region spanned by Bradi3g16331 and Bradi3g16320 may therefore be considered as a candidate for rust resistance.

**Figure 4:**
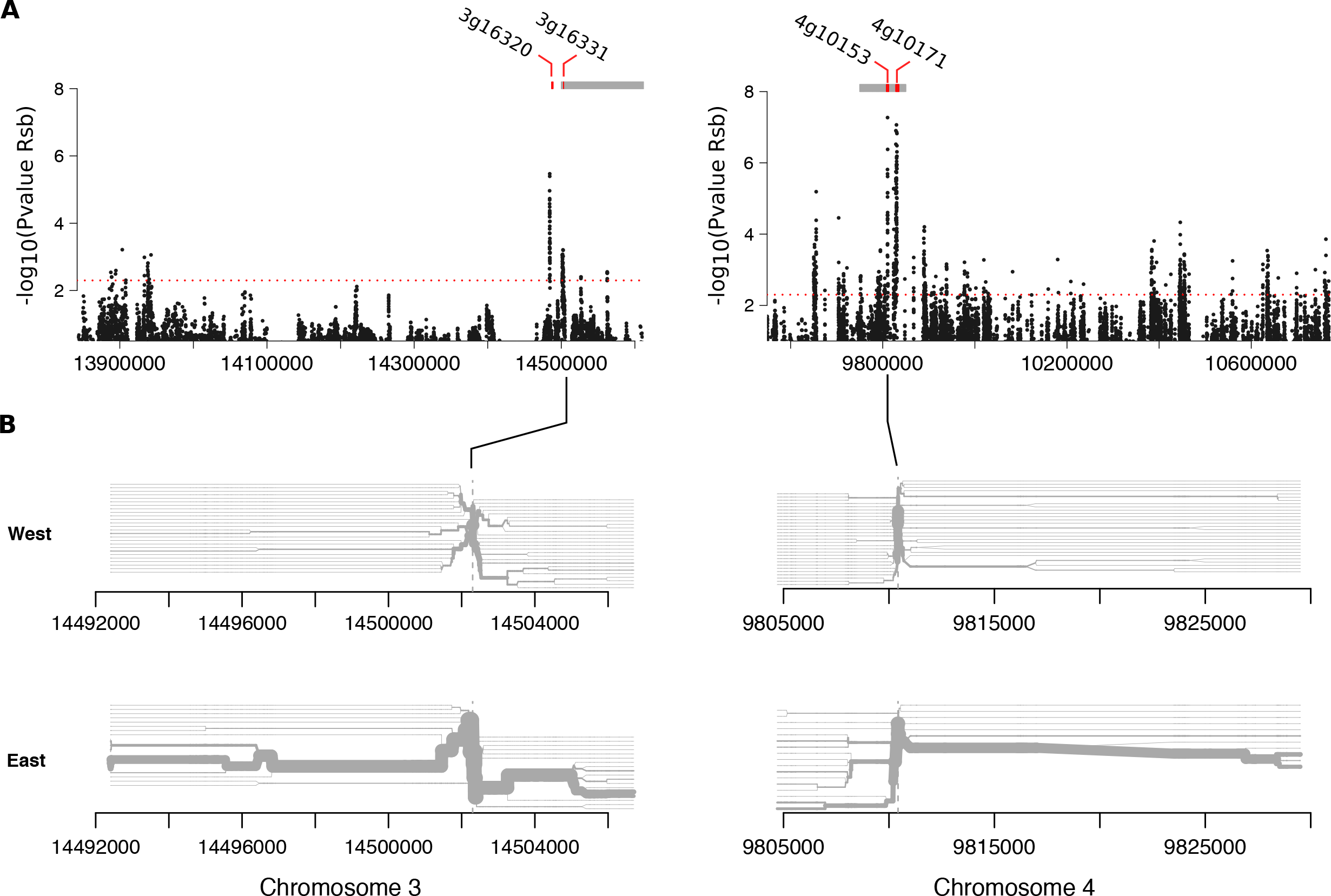
QTL involved in resistance to rust. A) Rsb signals observed in QTL_rust-3_ (left panel) and QTL_rust-4_ (right panel). The dashed line indicates the 1% −log_10_(Rsb P-values) outlier threshold. Note that in order to reduce the size of the figure, points with −log_10_(Rsb P-values) values <1 are not displayed. Grey boxes display the position of regions detected by diploS/HIC. Red bars display the position of genes detected by the Rsb test B) Haplotype bifurcation diagram in QTL sub-regions displaying strong of selection. Left panel: zoom in on chromosome 3; right panel: zoom in on chromosome 4. The diagrams visualize the breakdown of LD at increasing distances from the selected focal SNP, displayed by a vertical black line. Each horizontal line represents a haplotype. Each accession is represented by two lines, one for each haplotype. Horizontal blue lines are merged when two accessions share the same haplotype. The thickness of the line is therefore correlated to the number of accessions sharing the same haplotype.

The two adjacent Rsb signals in QTL_rust-4_ reach their highest point into two R-genes (Bradi4g10153: 9,807,879-9,812,927; and Bradi4g10171: 9,828,236-9,835,003; Figure 4A, right panel). These two genes are located in the near vicinity of the QTL peak (estimated to be located at 9,806,000 kb) and both belong to the eastern Rsb-diploS/HIC high-confidence gene set (Figure 4A, right panel). Bradi4g10153 and Bradi4g10171 constitute therefore further prime candidates for rust resistance. Congruent with the signal detected by the Rsb test, we observed extended haplotypes in the eastern population in both QTL_rust-3_ and QTL_rust-4_ (Figure 4B). The large majority of the eastern accessions display the extended haplotype in these regions, indicating a nearly completed selective sweep, especially in QTL_rust-4_ (Figure 4B).

QTL_rust-4_ shows a striking enrichment for genes involved in defense response (P-value = 8E-09) and in immune signaling processes such as phosphorylation (P-value = 5.6E-06). Among the 113 genes covered by QTL rust-_4_, 40 correspond either to a gene with a NBS-LRR, a receptor-like protein kinase (RLK) or a F-box domains, three types of genes that can confer resistance in plants. The presence of such a large gene clusters, as found in other regions of the genome (Table S2), further demonstrates the importance of narrowing down candidate regions under selection to top outlier genes for unbiased GO annotation.

## Discussion

Assessing the time and spatial scales at which selection acts is a key to understand how genetic diversity is maintained or lost through adaptation (Stinchcombe and Hoekstra 2008; Fuller et al. 2015). In plants, and especially in grasses, this question has been largely restricted to crops, which biases our understanding of evolutionary processes that have shaped genomes in their natural ancestors and extant relatives. In this study, we investigated the selective forces influencing adaptation in two populations consisting of 44 natural accessions of the wild Mediterranean grass *B. distachyon*. We found that ancient balancing and recent positive selection left distinct signatures on specific gene categories, and that positive selection affects distinct loci across the two populations. Importantly, our results support a role for pathogens in driving population differentiation and confirm that GWSS constitute effective approaches to pinpoint candidate genes as a complement to classical trait mapping.

### Time- and space-varying selection is shaping diversity in B. distachyon

The ending of the last glaciation period 10,000 years ago led to drastic and recent changes of plant communities in Eurasia (Svenning et al. 2008; Binney et al. 2017). At that time, climate warmed and species expanded over Europe (Hewitt 1999). Pollen-based studies show that vegetation expansion was fast, reaching up to 2km per year for some species (Hewitt 1999). To our knowledge, no fossil pollen records are available for *B. distachyon*, which prevents reconstructing the geographical distribution of this species before and during the last ice age. Yet a previous study showed that the two populations analyzed here experienced population size reduction during the last glaciation followed by a rapid expansion within the last 10,000 years (Stritt *et al.*, 2018). Even though unraveling the history of populations in southern peninsulas is more complex than in northern regions (Hewitt 2000; Feliner 2011), these results are congruent with the recent global postglacial recolonization of Europe by plants (Hewitt 1999; Svenning et al. 2008; Binney et al. 2017) and imply that *B. distachyon* populations had to adapt to newly colonized habitats in the recent past.

Balancing selection may have maintained ancestral polymorphisms over long periods of time in natural populations of *B. distachyon*. As a support of balancing selection, we identified an old set of shared polymorphisms with a coalescence approach. *B. distachyon* populations encounter heterogeneous habitats (Lopez-Alvarez *et al.*, 2015) which could have provided selective pressure strong enough to maintain polymorphisms over long periods of time within each population. Whether balancing selection results from spatial heterogeneity or from negative frequency-dependent selection remains yet to be investigated. Our results nevertheless suggest that adaptation to climate after recolonization does not necessarily involve *de novo* mutations, even after bottlenecks.

On the other hand, we also found evidence for positive selection acting on younger polymorphisms. Our analysis also revealed that positive selection is targeting different loci in the two populations and that these loci appear to be non-randomly distributed along chromosomes. As recombination rate is relatively high in *B. distachyon* (Huo et al. 2011) and since we observed a rather moderate linkage disequilibrium in the two populations, we do not believe that this pattern is due to extended linkage disequilibrium and the subsequent process of background selection along large genomic regions (Cutter and Payseur 2013; Slotte 2014). Rather, the peaks of selection we identified were narrow and allowed to pinpoint genes, indicating that while *B. distachyon* is predominantly inbreeding, outcrossing events must occur every now and then to limit extended linkage disequilibrium. This is also congruent with the slightly higher level of heterozygosity we observed here in *B. distachyon* compared to other selfing plants such as *A. thaliana* (Platt et al. 2010). Interestingly, the regions displaying signs of soft-sweep or ongoing selection tended to be larger in the western population than in the one of the eastern cluster despite slower LD decay in this latter group. This also indicates that selection may have acted more recently in the western population, possibly due to the modern and rapid expansion of these lineages in the Spanish peninsula (Stritt *et al.* 2018).

*B. distachyon* occurs exclusively around the Mediterranean rim which constitute a mosaic of landscapes. Our results reveal that this variation in habitats lead to natural selection which affected different genes and genomic regions across populations. As we obtained enrichments for different GO terms in the eastern and western populations, we also imply that contrasting abiotic and biotic factors are shaping natural population diversity in different regions of the genome of *B. distachyon* through positive selection. Even though we identified more genes under positive than under balancing selection, it would be daring to conclude that the former selection regime, less challenging to detect (Delph & Kelly, 2014), is a predominant process shaping diversity in this species. Rather, we believe that we provide here genomic evidence that balancing selection also leads to the adaptation of *B. distachyon* populations and that selection acted at different temporal scale.

### Pathogens as a potential driving force of population evolution

Host-pathogen interactions lead to a strong coevolutionary dynamics and are considered a major factor shaping diversity (Fumagalli et al. 2011; Karasov et al. 2014; Krattinger and Keller 2016a). Two main types of interaction have been proposed. Under an arms race model, repeated innovation from both sides results in repeated fixation of advantageous alleles (Brown and Tellier 2011). This interaction can therefore lead to positive selection that can be detected by tests focusing on extended haplotypes. The other type of interaction is often referred to as Red Queen dynamics or trench warfare, where alleles involved in the interaction are recycled through negative-frequency dependence and can therefore subsist for long periods of time in populations through balancing selection (Brown and Tellier 2011).

The plant immune system machinery is complex and is composed of two tiers of extracellular and intracellular receptors (Krattinger and Keller 2016a; Krattinger and Keller 2016b) that efficiently detect the presence of pathogens (for review Greeff *et al.*, 2012; Couto & Zipfel, 2016; Eckardt, 2017). In this study, we found a significant enrichment of signals of selection at genes involved in this first level of defense in the eastern population. More specifically, we found many of well-characterized R-genes, i.e. genes with NBS-LRR domains and involved in disease resistance (Mchale *et al.*, 2006; Jacob *et al.*, 2013; Liu *et al.*, 2014; Couto & Zipfel, 2016; Eckardt, 2017; Ooijen *et al.*, 2017), to be under ongoing or completed positive selection in the eastern population. On the other hand, we also found additional R-genes and genes involved in phosphorylation, a process especially important for immune signaling response in plants (for review Park *et al.*, 2012), to be under balancing selection. Our results are thus consistent with the two classical models of host-pathogens coevolution (Mondragón-palomino et al. 2002; Mchale et al. 2006; Gos et al. 2012; Mace et al. 2014; Zhong et al. 2015; Wu et al. 2017). Overall, and as shown in other organisms (Karasov *et al.* 2014; Fumagalli *et al.*, 2011; Krattinger & Keller, 2016a,b; Bourgeois *et al.*, 2017), our genome-wide approach suggests that pathogens may constitute an important driving force of population and genome evolution in *B. distachyon*.

The significant enrichments for R-genes and for genes associated to the process of defense response in the gene sets under positive selection were only observed in the eastern population. As we incorporated the demographic history to our analyses, we conclude that the bottlenecks experienced by both populations did not extensively bias our results and did not interfere with our conclusions on the importance of host-pathogen interaction in the eastern population. The geographical origin of the populations could, however, play a role in this observation. The Middle East is indeed the center of origin of many grasses, including *B. distachyon*, and of their associated pathogens (Wyand & Brown, 2003; Stukenbrock *et al.*, 2005; Opanowicz *et al.*, 2008; Hovmøller *et al.*, 2011). Many studies found that both resistance genes in plants and effector genes in pathogens can be organized in clusters evolving through an arm race and resulting in a gene birth-and-death process (Michelmore and Meyers 1998; Dong et al. 2015; Singh et al. 2015). Because centers of origin are usually associated to higher diversity, it is therefore possible that a higher level of pathogen diversity drove selection at a larger number of resistance genes in the eastern population.

### GWSS as complementary approach to QTL and GWAS

Disentangling the mechanisms that promote or prevent adaptation requires more integrated studies, using both experimentations in controlled conditions and methods to characterize genetic diversity in natural populations (Feder and Mitchell-olds 2003; Stinchcombe and Hoekstra 2008; Flood and Hancock 2017). Several studies used GWSS to validate genes functionally characterized or previously identified through GWAS and to propose stronger hypotheses on the mode of selection operating on traits relevant for adaptation (Roulin *et al.* 2016; Tang *et al.*, 2007; Fumagalli *et al.*, 2011; Bourgeois *et al.* 2017). Following this idea, we inspected three QTL regions responsible for the resistance of *B. distachyon* to the leaf-rust fungus *Puccinia brachypodii*, a natural pathogen of *B. distachyon* expected to exert selection on natural accessions.

For two of these QTL regions on chromosomes 3 and 4, we found signs of positive selection and reduced haplotype diversity in the eastern cluster in the vicinity of the QTL peaks, but no sign of balancing selection. These results strongly indicate that positive selection is shaping rust resistance in *B. distachyon* natural populations, as suggested in other species (Dodds & Thrall, 2009; Chavan *et al.*, 2015). On chromosome 3, our analysis identified a 16kb region spanned by an unknown protein and a serine/threonine phosphatase, a class of genes known for their role in defense response and stress signaling (País et al. 2009; Durian et al. 2016), to be putatively involved in resistance to rust. The region identified on chromosome 4 is more complex and consists of a cluster of resistance and stress signaling genes. Nevertheless, and while other genes display evidence of positive selection, two R-genes that belong to the eastern high-confidence gene set co-localize with the peak of the QTL. Such genes have been shown to confer resistance to rust in other species (Bettgenhaeuser et al. 2014) and constitute further prime candidates for functional characterization.

*B. distachyon* is closely related to major crop cereals as well as to grass species used for biofuel production. Translating research from *B. distachyon* to plants of agronomical and economical interest will require a deeper understanding of the genetic architecture of traits involved in the response to environmental stresses. The molecular basis of tolerance to various abiotic stresses such as drought, salt and cold has been investigated in this species (Luo et al. 2011; Manzaneda 2013; Carmo and Charron 2014; Gordon et al. 2014; DesMarais and Juenger 2015; Sun et al. 2015; Mur and Bosch 2016). Here, we also highlighted cadmium pollution as a potential factor of positive selection in the eastern population. As pollution with heavy metals including cadmium has been reported in Turkey in regions where accessions were collected for this study (Bakirdere and Yaman 2008; Mor and Ceylan 2017), our results suggest that *B. distachyon* could be used to investigate the tolerance to this stress. As genetic transformation is highly efficient in this species relative to other grasses, we anticipate that combining classical trait mapping analyses with GWSS will assist allele mining for additional eco-responsive traits.

### Conclusion

Our results revealed widespread signatures of natural selection at genes involved in adaptation in *B. distachyon* and provide the community with a list of candidate genes displaying strong signs of selection in natural populations. We also found that pathogens may constitute an important driving force of genetic diversity and evolution in this system. While we limited our analysis to classical point mutations, recent studies showed that copy number variants (CNVs) and transposable element polymorphisms are abundant across *B. distachyon* populations (Gordon *et al.*, 2017; Stritt *et al.*, 2018). Hence, the important genomic resources currently developed in this species open new avenues of research to further investigate the role of structural variation in natural population evolution and adaptation. To date, *B. distachyon* remains a classical model for research on grass genomics with a strong orientation towards applied research. Thanks to the high quality of its reference genome and the existence of large collections of freely available natural accessions collected from the species native range, it also constitutes a prime system for studies in ecology, population genomics and evolutionary biology.

## Experimental procedures

### SNP calling, population structure and genetic diversity

We used paired-end Illumina sequencing data generated for 44 accessions of *B. distachyon* (Gordon *et al.*, 2017; Table S1 for information about the origin of the accessions and sequencing effort) originating from Spain (N=16), France (N=1), Turkey (N=23) and Iraq (N=4). Reads were aligned to the reference genome v2.0 with BWA-MEM (standard settings; Li, 2013). After removing duplicates with Sambamba (Tarasov et al. 2017), single nucleotide polymorphisms (SNPs) were called with Freebayes (Garrison and Marth 2016). The output was then filtered by removing SNPs with more than 10 missing genotypes or more than 2 alleles, a quality lower than 20 and a mean depth lower than 20 or higher than 200. Data were phased using the software BEAGLE V4 (Browning and Browning 2007) using default settings.

We then used the program Admixture (Alexander and Novembre 2009) to identify the genetic structure of the two populations. The analysis was run for K values from 1 to 5, and the best model was determined as the model with the lowest cross-validation error. Summary statistics such as within-population nucleotide diversity (π) were computed with the R package PopGenome (Pfeifer et al. 2014). Levels of heterozygosity were calculated with VCFtools (Danecek et al. 2011).

### Detecting positive selection associated to nearly completed sweeps

We used the Rsb test (Tang et al. 2007) to detect signatures of recent or almost completed hard sweeps. This test detects haplotypes that are positively selected in one population by estimating the length of haplotypes around each allele at a core SNP and then comparing these lengths between populations. While the output of the test provides P-value of significance, it also indicates in which population a given allele is under selection. Rsb statistics were computed for each SNP with the R package rehh2.0 (Gautier et al. 2012) with default settings.

### Detecting ongoing positive selection and soft-sweeps within populations

We used the software H-scan (Schlamp et al. 2016) to detect soft-sweep or loci under incomplete ongoing positive selection. To do so, we calculated average pairwise haplotypes lengths using the number of segregating sites spanned by each tract within each population. The H-scan statistics is expected to be larger as the number of extended haplotypes increases in a population. We ran the method on eastern and western accessions independently to detect soft-or incomplete selective sweeps within each geographical group.

### Testing for functional clustering and subsequent filtering of GWWS results

We first performed a GO annotation for the 32,712 genes annotated in the reference genome (version 2.1, https://phytozome.jgi.doe.gov) with Blast2GO (Conesa et al. 2005). We then controlled for potential gene clustering by following the procedure described in (Al-Shahrour *et al.*, 2010). The entire gene set of the reference genome was split into windows of 50 consecutive genes. Windows were moved along chromosomes in steps of 25 genes to allow for half-window overlaps. Enrichment analyses of biological processes were then performed for all the generated windows with the R package GOstats (Falcon and Gentleman 2017) using Fisher’s exact test. P-values were subsequently adjusted for multiple testing with a Benjamin-Hochberg correction. Regions were considered significantly enriched for a biological process when they displayed a corrected P-value ≤ 0.01 and also harbored at least five genes associated to the given process.

Both the H-scan and the Rsb tests compute statistics at each SNP. To limit false positives, we first selected 10 kb windows displaying at least four significant SNPs within the top 1% outliers. Overlapping significant windows were merged. To limit the impact of gene clustering on the GO analysis, we then narrowed down the selected windows by keeping only the genes located at and around (−10% of the peak value) each of the top 1% peaks of selection. These filtering criteria, however, were not applied to the outputs of ARGweaver and diploS/HIC, which are window-based approaches.

### Demographic history and filtering false positives

We used the recently developed machine-learning algorithm diploS/HIC (Kern & Schrider, 2018) to further control for the potentially confounding effects of demography on our outlier approach. To do so, genomic windows were simulated under neutrality, with a hard sweep or with a soft sweep. We trained the algorithm using a set of 3,000 coalescent simulations obtained with discoal (Kern and Schrider 2016). We simulated 330kb windows each divided into 22 subwindows. Hard and soft sweep examples consisted in windows with a sweep occurring in the two central 30kb subwindows. The adjacent subwindows are assigned as linked to selection.

We performed simulations for each of the two populations using demographic trajectories inferred in a previous study (Stritt et al. 2018). Note that we rescaled these estimates with a mutation rate of 1.4×10-9/bp/generation instead of the previously used 7×10^−9^/bp/generation. We estimated this new mutation rate by aligning the orthologs of 100 genes and using rice as an out-group (divergence estimated at 40My; The International Brachypodium Consortium, 2010). We sampled selection coefficients from an uniform prior 2Ns ~ (250,25000) for the eastern population and 2Ns ~ (2000,200000) for the western population, to take into account the largest estimate for present effective population size (N) in the western cluster (N≈4,000,000) compared to the eastern one (N≈500,000). We used a truncated exponential prior for recombination rates encompassing the range 5.9×10^−9^−1.18×10^−7^, values that are between ten times lower and two times higher the rate of 5.9×10^−8^ estimated from controlled crosses (Huo et al. 2011).

To take into account the likely heterogeneity in effective population sizes and mutation rates along the genome, we used priors for present effective population sizes between twice lower and five times higher than the values estimated from the SMC++ analysis for present times (Stritt et al. 2018). We conditioned on sweep completion occurring between present time and shortly before populations expansion, around 10,000 generations ago. For soft sweeps, we used uniform priors on the initial frequency of the adaptive variant of (0,0.2). We took into account inbreeding by assuming each individual contained two copies of the same haplotypes.

This set of simulated datasets was then used to train a supervised machine-learning algorithm to differentiate between each category. Predictions on the actual genomic dataset were then performed over subwindows. Since the statistics computed by diploS/HIC are normalized across subwindows, they can remain flat if selection is strong and erases diversity over the whole region to be classified. We therefore classified genomic subwindows as selected, linked or neutral using two sizes (15 and 50kb), in order to detect genomic regions displaying the strongest signals of selection. The algorithm as well as a detailed tutorial are available at https://github.com/kern-lab/diploSHIC.

We finally used bedtools intersect (Quinlan and Hall 2010) to identify high-confidence regions recovered by both the Rsb test and diploS/HIC or by both the H-scan test and diploS/HIC. Genes present in such high-confidence regions were retrieved with bedtools intersect using the annotation of the genome of *B. distachyon* v2.1 (https://phytozome.jgi.doe.gov).

### Distribution of genes under positive selection along chromosomes

To compare the distribution along chromosomes of candidate genes for recent positive selection (DiploS/HIC-Hscan and DiploS/HIC Rsb candidate genes combined) in each population, we used linear models where the density of selected genes along each chromosome identified in the eastern and western populations (100,000 bins per chromosome) were entered as variables. We eventually used circos (Krzywinski et al. 2009) to visualize the density of genes under positive selection along each chromosome.

### Detecting ancient balancing selection with ancestral recombination graphs (ARG)

We used the software ARGWeaver to detect additional candidate regions for balancing selection with a coalescence approach (Rasmussen et al. 2014). We included in the analysis a subset of 12 accessions with high sequencing depth and covering the largest geographical range (6 accessions from each population) to limit computation time. We used a mutation rate of 1.4×10-9/bp/generation and a recombination rate of 5.9×10-8/bp/generation. Note that we subsequently used an outlier approach to identify the oldest polymorphisms present in the two populations (top 1% outliers). Therefore, potential biases inherent to the use of a molecular clock do not affect our analysis. ARGWeaver is also flexible with regard to recombination rate as it reconstructs ancestral recombination graphs and accommodates variable recombination rates, effective population sizes and genealogies along the genome. The algorithm was run for 1000 iterations, using 20 discretized time steps, a maximum coalescence time of 3 million generations and a prior effective population size of 100,000 individuals. This size was of the same order of magnitude as the harmonic mean of effective population sizes over the last 500,000 generations (spanning the recent population declines and expansions), as estimated in a previous study (Stritt et al. 2018) using the algorithm SMC++ (Terhorst et al. 2017).

### GO annotation of genes under selection

For each high-confidence gene set and the gene set under balancing selection, we extracted the genes located in the filtered regions with bedtools (Quinlan and Hall 2010). We then examined potential enrichment for biological processes for each of the selected gene sets with the R package GOstats (Falcon and Gentleman 2017) using the Fisher’s exact test. Gene sets were considered significantly enriched for a biological process when they displayed a P-value ≤ 0.01 and harbored at least five genes associated to the given process. The ancestor and child terms of each significant process were determined using QuickGO (http://www.ebi.ac.uk/QuickGO) and used to simplify Fisher test outputs and keep non-redundant terms.

We also retrieved from each gene set under selection the number of genes containing a NBS-LRR domain, and annotated as R-genes in version 2.1 of the genome. We then estimated whether these gene sets were enriched for R-genes compared to the rest of the genome with Fisher’s exact test in R (R development core team, 2017).

### QTL for leaf-rust resistance validation

A QTL analysis performed in *B. distachyon* revealed three genomic regions involved in the resistance to *P. brachypodii* (Barbieri et al. 2012). The coordinates of these three QTL were extracted from (Barbieri et al. 2012) from v.2.1 of the *B. distachyon* reference genome. We then assessed weather those three regions were identified as outliers in at least one of the tests of selection or in the high-confidence gene set. We further visualized the extension of haplotypes at candidate regions using the bifurcation.diagram() function of the rehh2.0 R package.

## Acknowledgements

We thank the Genetic Diversity Center-ETH Zurich for providing access the Euler high-performance cluster. This research was carried out on the High Performance Computing resources at New York University Abu Dhabi and ETH-Zurich. We also would like to thank Beat Keller’s group as well as Mahendra Mariadassou for their advices during the elaboration of the study. This work is supported by the Ambizione program of the Swiss National Science Foundation (PZ00P3_154724). The work conducted by the U.S. Department of Energy Joint Genome Institute, a DOE Office of Science User Facility, is supported under Contract No. DE-AC02-05CH11231.

## Availability of data and materials

The raw vcf file and the raw outputs of each GWWS will be archive upon acceptance of the manuscript. All whole-genome sequences data are available at the NCBI Sequence Read archive (SRA available in Gordon *et al.*, 2017).

## Supporting Information

**Figure S1: Admixture analysis**

Cross-validation plot.

**Figure S2: Confusion matrices for diploS/HIC.**

The trained algorithm was applied on datasets simulated under a coalescent framework, and the proportion of simulations correctly assigned to their category was estimated. The location of the classified subwindows relative to the sweep is shown vertically and the predicted class for each subwindow is shown horizontally. Each row corresponds to 3,000 simulations.

**Table S1:** Geographical coordinates of the 44 accessions and sequencing effort.

**Table S2:** Significant GO terms in each 50 gene-window of the reference genome.

**Table S3:** List of genes under selection.

